# Multiple Causal Variants Underlie Genetic Associations in Humans

**DOI:** 10.1101/2021.05.24.445471

**Authors:** Nathan S. Abell, Marianne K. DeGorter, Michael Gloudemans, Emily Greenwald, Kevin S. Smith, Zihuai He, Stephen B. Montgomery

**Affiliations:** Department of Genetics, School of Medicine, Stanford University, Stanford, CA, 94305, USA; Department of Pathology, School of Medicine, Stanford University, Stanford, CA, 94305, USA; Biomedical Informatics Program, Stanford University, Stanford, CA, 94305, USA; Department of Neurology and Neurological Sciences, Stanford University, Stanford, CA 94305, USA; Quantitative Sciences Unit, Department of Medicine, Stanford University, Stanford, CA, 94305, USA

## Abstract

The majority of associations between genetic variation and human traits and diseases are non-coding and in strong linkage disequilibrium (LD) with surrounding genetic variation. In these cases, a single causal variant is often assumed to underlie the association, however no systematic assessment of the number of causal variants has been performed. In this study, we applied a massively parallel reporter assay (MPRA) in lymphoblastoid cells to functionally evaluate 49,256 allelic pairs, representing 30,893 genetic variants in high, local linkage disequilibrium for 744 independent cis-expression quantitative trait loci (eQTL) and assessed each for colocalization across 114 traits. We identified 8,502 allele-independent regulatory regions containing 1,264 allele-specific regulatory variants, and found that 17.7% of eQTL contained more than one significant allelic effect. We show that detected regulatory variants are highly and specifically enriched for activating chromatin structures and allelic transcription factor binding, for which ETS-domain family members are a large driver. Integration of MPRA profiles with eQTL/complex trait colocalizations identified causal variant sets for associations with blood cell measurements, Asthma, Multiple Sclerosis, Inflammatory Bowel Disease, and Crohn’s Disease. These results demonstrate that a sizable number of association signals are manifest through multiple, tightly-linked causal variants requiring high-throughput functional assays for fine-mapping.

## INTRODUCTION

Genome-wide association studies (GWAS) have emerged as an important tool to assess the effect of individual genetic variants on phenotypes ranging from gene expression to complex traits and diseases (*1, 2*). However, due to linkage disequilibrium (LD), it is often not possible to identify a single highly-associated and likely causal genetic variant from multiple correlated variants. Furthermore, it is typically assumed that only a small fraction of variants or a single variant will causally explain a GWAS significant locus. To address this challenge, statistical and functional fine-mapping approaches have been developed to identify credible sets of variants most likely to harbor the causal allele (*3–7*). However, these techniques often cannot distinguish between extremely proximal or highly-linked variants and lack systematic prior information on the number of causal variants underlying association signals.

One approach to systematically identify causal variants while controlling for the confounding effects of LD is through massively parallel reporter assays (MPRAs). MPRAs measure the effects of synthetic DNA libraries on expression of a reporter gene, like luciferase or GFP, containing a 3’ UTR barcode (*8*). They have been used to screen potential regulatory elements in diverse cellular contexts, as well as for saturation mutagenesis or tiling along individual regulatory regions of interest (*9–13*). Beyond regulatory function, MPRAs have also been increasingly applied to assay the differential effects of allelic sequence pairs representing natural human variation, focused on variants that are human-specific or are lead variants of signals in GWAS or expression quantitative trait loci (eQTL) (*14–16*). However, despite increases in throughput no prior work has systematically dissected linked haplotype blocks in humans to assess the distribution of causal alleles; previous studies either targeted only variants with the strongest associative p-values with respect to particular phenotypes and/or applied extensive prior filtering to reduce library size for sensitive cell types (*14, 15, 17, 18*). In *S. cerevisiae*, high-resolution QTL mapping enabled by inbred crossing and biochemical phenotypic characterization has identified many instances of multiple linked causal variants influencing the same traits, suggesting that the same may occur in humans (*19, 20*).

In this study, we applied an MPRA to systematically characterize the distribution of causal variants underneath a large number of eQTL and GWAS loci. We identified regulatory variation from a pool of 49,256 linked genetic variants, most of which were strongly associated with gene expression in lymphoblastoid cell lines (LCLs). We found 8,502 regions with significant allele-independent regulatory effects and 1,264 variants with significant allele-dependent effects. We showed that variants with significant MPRA effects are highly enriched for diverse signatures of regulatory activity including allelic imbalance in transcription factor binding and chromatin accessibility. Of the 744 genes regulated by these variants as eQTL, 17.7% were the target of more than one significant allelic MPRA effect with many in high LD. We also identified dozens of GWAS-eQTL colocalizations with extensive MPRA coverage, including causal variant sets underlying joint associations with *ORMDL3*/Asthma, *AHI1*/Multiple Sclerosis, *PACSIN2*/Platelet Count and *ERAP2*/Crohn’s Disease/Inflammatory Bowel Disease. These results provide new evidence for specific variants underlying hundreds of independent genetic associations, including both gene expression and complex traits, as well as highlight a notable fraction of associations caused by the allele-dependent action of multiple variants.

## RESULTS

### Functional fine-mapping of eQTL reproducibly identifies regulatory and allelic hits

We selected independent, top-ranked eQTL signals across 744 eGenes identified in the CEU (Utah residents with Northern and Western European ancestry) characterized as part of the HapMap (*21*). Each eQTL had a median of 6 top associated variants (range 1-472) in perfect LD. For each lead variant, we identified all additional variants with r^2^ >= 0.85 that were at least nominally associated with the same gene. Our final library included 30,893 variants, with a median of 50 variants per eQTL locus (range 2-2824) all of which were highly associated with gene expression and were in high LD with their respective lead variant (Figure 1A). For each variant, we extracted 150 bp genome sequences (centered on the variant) and generated a MPRA library by random barcoding (Figure 1B). In 150 bp windows with more than one variant, distinct oligos were designed for each possible haplotype resulting in an average of 3.19 designed oligos per variant and 49,256 total allelic pairs. After reporter gene insertion, the library was transfected into LCLs in triplicate, sequenced and then quantified for each oligo (Methods).

**Figure 1 -.**
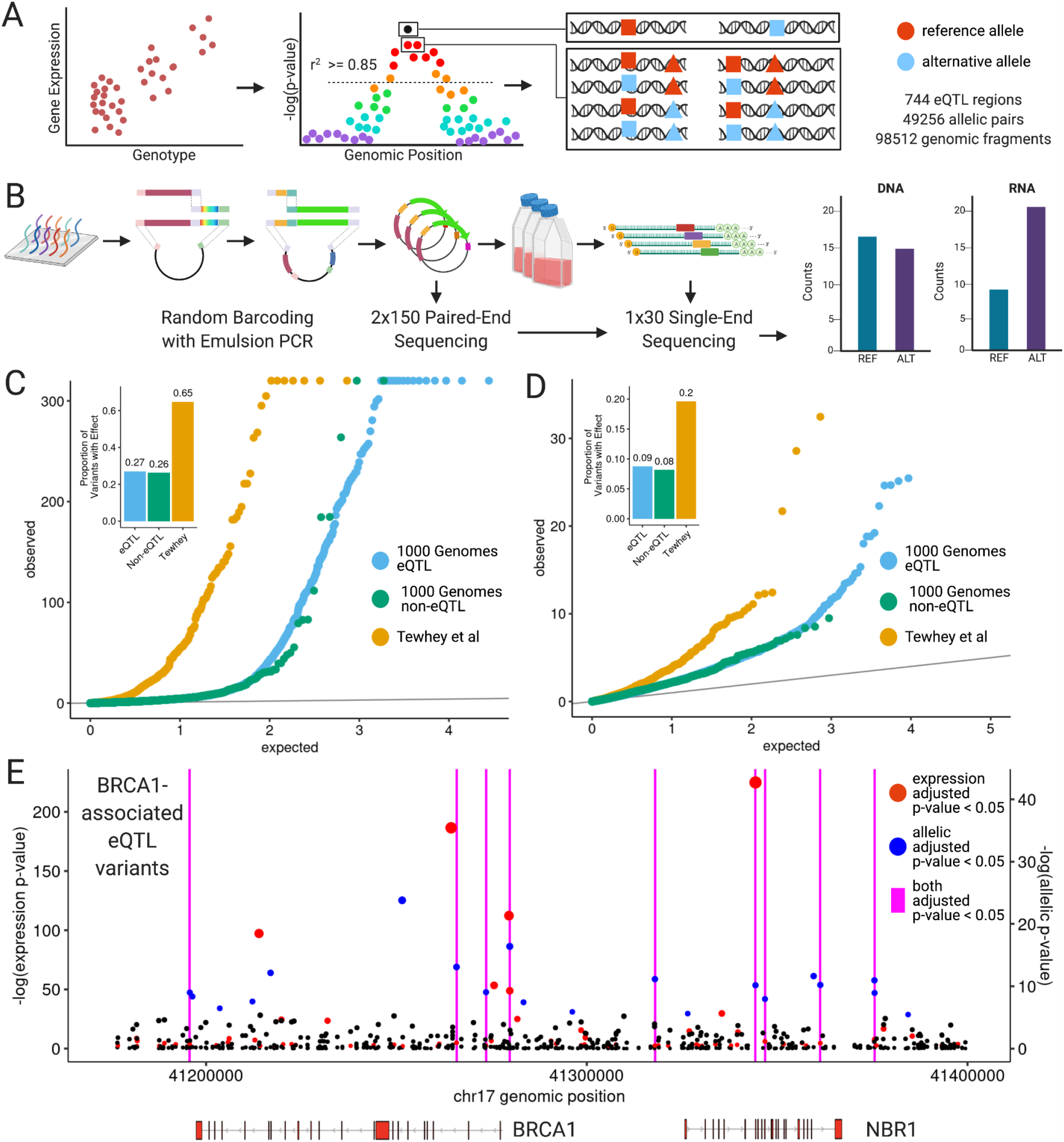
Design and implementation of a linked variant massively parallel reporter assay. (A) Variant selection and oligonucleotide sequence design of a variant-focused MPRA. (B) Experimental procedure random barcoding and expression of the MPRA library in cultured human lymphoblastoid cells. (C) Distribution of eQTL, non-eQTL, and (*14*) variant expression p-values and relative effect proportions. (D) Same as in (C) but with allelic p-values. (E) Genomic position and unadjusted p-values for all tested BRCA1-associated variants with colors indicating adjusted p-value <= 0.05. Vertical magenta lines indicate positions of variants where both effects are significant.

To infer regulatory effects from oligo counts, we used negative binomial regression as implemented in DESeq2 using the approach outlined in (*22*). We computed two sets of summary statistics to test, first, the allele-independent regulatory effects of an oligo (“expression” effects) and second, the difference in regulatory effects between reference and alternative oligos of the same variant (“allelic” effects). We detected 8,502 expression effects and 1,264 allelic effects across all tested variants. We observed a small increase in the total number of significant effects in eQTL relative to non-eQTL, possibly reflecting the low proportion of significant associations which are expected to be causally relevant (Figure 1C, 1D). However, we observed a larger increase in effect size among significant variants in eQTL relative to non-eQTL that was absent in non-significant effects (Supplementary Figure 1D). Taken together, we obtained for each eGene a profile of allele-independent and -dependent effects across all highly-associated variants in the local cis-window (Figure 1E).

By design, a subset of our variants (N=782) were previously identified as expression-modulating variants in (*14*). To our knowledge, no previous studies have systematically replicated MPRA effects to this extent, and it is notable that this overlapping subset contained a highly elevated number of both expression and allelic effects relative to other tested variants (Figure 1C, 1D). Our assay also included a set of additional variants from (*14*) that were not significant in that study, as well as some additional coincidental overlap. We observed that 89.6% of variants with significant effects in both datasets were directionally concordant while ∼62% and ∼50% of variants with significant effects in one or neither dataset respectively were concordant (Supplementary Figure S2A). Based on these results, we constructed a concordant, high-confidence “MPRA positive” variant set containing 250 variants with expression effects and 120 with allelic effects that are highly consistent between both data sets (Supplementary Figure S2B, S2C).

### Diverse transcription factor programs contribute at eQTL

The large number of MPRA expression effects provided an opportunity to identify transcription factors impacting gene expression within eQTL. We observed widespread enrichment of ChIP-seq peaks for multiple transcription factors in MPRA expression effects (N=160 total TFs) (*23, 24*). These enrichments were exclusively positive, indicating increased representation in significant versus non-significant expression effects. Moreover, applying a more stringent filter for expression effects (adjusted p-value <= 5e-10) substantially increases these enrichments in most transcription factors with odds ratios ranging from 1.2- to 17-fold (Figure 2A, Supplementary Table S5). This demonstrates the wide range of regulatory element effects captured in our assay and highlights the role of multiple transcription factors in driving the regulatory effects of genetic variation.

**Figure 2 -.**
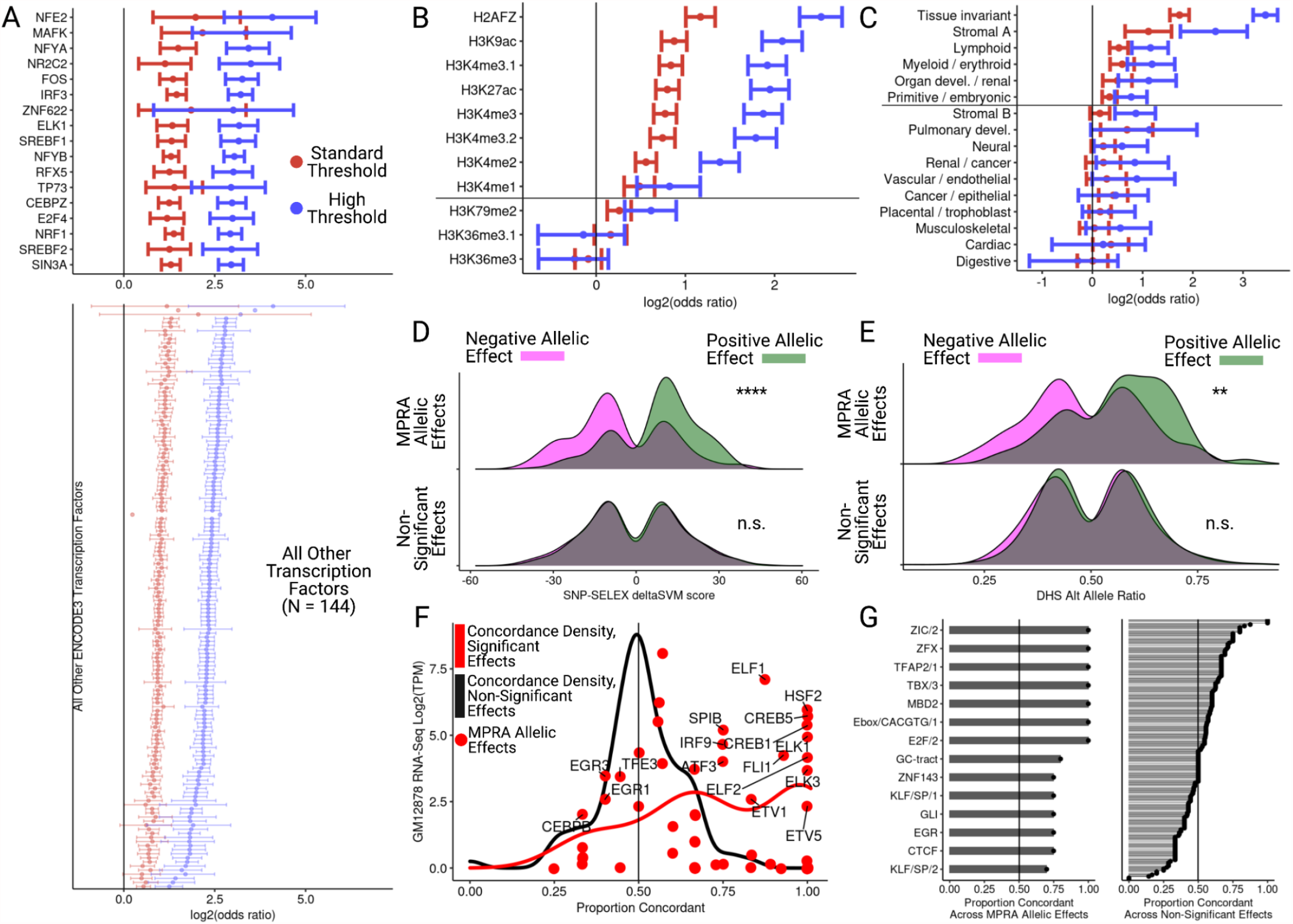
General and allele-specific functional properties of regulatory variants. (A) Odds ratios and 95% confidence intervals for enrichment of ENCODE ChIP-seq peaks in fragments with expression effects. Only transcription factors with an enrichment adjusted p-value < 0.005 are shown. (B) Same as in (A) but for histone modifications and all tested marks are shown; marks above the horizontal line have an adjusted p-value < 0.05 at both thresholds. (C) Same as in (A) but for clustered chromatin accessibility regions in fragments with expression effects. (D) Distribution of SNP-SELEX deltaSVM scores at allele-specific binding variants stratified by MPRA allelic effect direction (color) and significance category (top and bottom); MPRA allelic effects have adjusted expression and allelic p-values <= 0.05 while non-significant variants have p-values > 0.75. (E) Same as in (D) except for allelic imbalance in chromatin accessibility from ENCODE. (F) For all TFs evaluated in (D), comparison of the concordance proportion across MPRA variants with the expression of each included TF in GM12878 cells; curves indicate the concordance densities on the x-axis for significant and non-significant MPRA allelic effects with points indicating only significant effect concordances. (G) Comparison of directional concordances within accessible chromatin motifs for significant (left) and non-significant (right) MPRA effects. * <0.05, **<0.005, ***<0.0005, ****<5e-5.s

We next evaluated all available histone modifications included (*23*) and observed enrichments for activating histone modifications but not for repressive marks like H3K36me3 (Figure 2B). Notably, as our assay was performed using a non-integrated plasmid library, these effects indicate that genome sequence alone rather than downstream histone modifications are the principal drivers of expression changes induced at these loci. We observed the strongest enrichments in chromatin accessibility regions that were tissue invariant or specific to the Stromal A (representing JDP2 and other AP-1 transcription factor family motifs), Lymphoid, and Erythroid/Myeloid tissue clusters, demonstrating that cell-type information encoded in accessible chromatin patterns is detectable in our assay (Figure 2C) (*25*). Consistent with histone enrichments, this indicates that the effects of many transcription factors do not require chromatin remodeling activity *per se* to regulate transcription if the sequence is accessible.

### Identifying regulatory variants by allelic transcription factor binding and chromatin accessibility

Given that multiple transcription factors were enriched in expression hits at eQTL, we aimed to dissect the subset of transcription factors that were targets for regulatory variation by characterizing allelic MPRA hits. We first assessed whether the direction of allelic effect (i.e. higher expressing reference or alternative allele) was concordant with metrics of allele-specific activity in the same molecular contexts. We observed a global concordance between SNP-SELEX scores, a set of allele-specific binding models based on *in vitro* TF binding affinities, for variants with significant allelic effects (Fisher’s exact p-value = 3.43e-15) that was absent in variants with non-significant effects defined as those variants with both expression and allelic adjusted p-value >= 0.5 (Fisher’s exact p-value = 0.63, Figure 2D) (*26*). In order to identify the transcription factors driving these allelic effects, we computed the concordance proportion (i.e. how often a SNP-SELEX score for a specific TF was concordant with the MPRA allelic effect) for all transcription factors overlapping at least three tested variants (Figure 2E). The mean concordance proportion across TFs (N=59) was 0.733 when using significant MPRA variants, and was 0.505 when using the non-significant variants (N=91) (binomial logistic GLM p-value = 8.46e-12). In the two cases where we observed a decrease in expected concordance within significant MPRA variants, the TFs involved were generally lowly-expressed in GM12878 except for *EGR1/3* and *CEBPB*, which can function as both an activator and a repressor depending on cellular context (*27*). Notably, we also observed significant expression or allelic MPRA effects in 27.6% of all tested and significant SNP-SELEX variants. While substantial, this implies that a minority of expected allele-specific TF binding effects have direct transcriptional consequences without other contextual effectors, and that MPRAs provide a powerful tool to link allelic binding in particular with a functional outcome.

A similar pattern emerged when comparing allelic imbalance in accessible chromatin with MPRA allelic effects (*28*). We observed significant concordance between allelic imbalance and MPRA allelic effect directions for significant MPRA effects but not those with insignificant effects (Fisher’s exact p-value = 7.33e-3 and p-value = 0.839 respectively; Figure 2F). Separation by functional footprint within accessible chromatin regions revealed that several motifs, including Gli and a canonical E-box, were perfectly concordant across all tested variants with significant effects. In contrast, concordances derived from variants with non-significant effects were distributed around 0.5 (Figure 2G).

To further explore the relationship between regulatory variants and chromatin accessibility, we also integrated two sets of caQTL data and attempted to identify variant annotations which increase MPRA signal (*28, 29*). Using allelic imbalance data from ENCODE, we found that significant MPRA allelic effects were strongly concordant with allelic imbalances for variants located within their associated DHS peaks, but not variants located adjacent to them (Fisher’s exact p-value = 3.2e-5 and p-value = 0.055, respectively; Supplementary Figure S3A). Separately, in a set of caQTLs assessed across ten population groups, variants which were significant caQTLs in multiple populations were more enriched in MPRA allelic effects than variants which were significant caQTLs in only a few populations (Supplementary Figure S3B). Thus, variants which are located inside their associated accessible region and are replicated between populations not only offer more plausible biological mechanisms, but are also enriched for allelic MPRA effects despite the use of a non-integrated library. Taken together, MPRA allelic effects were significantly concordant with both *in vitro* and *in vivo* measures of allelic regulatory activity while variants with non-significant effects were directionally random.

### MRPAs inform advantages and limitations of integrative non-coding variant effect prediction

A major ongoing challenge is to summarize and predict the regulatory effect of non-coding variants using sequence and sequence annotation alone. We evaluated whether state-of-the-art genome-wide non-coding variant effect predictors could identify genetic variants with significant allelic effects detected by our MPRA. Using principal component scores from Enformer, a neural network that incorporates sequence information across 200kb windows, we observed significant enrichment of allelic variants in the top percentiles of Enformer scores (Figure 3A inset) (*30*). We next annotated all tested variants with their annotation principal components (aPCs) from FAVOR, an integrated variant effect prediction tool whose scores are computed across all variants in TOPMed (*31*). We again observed differences in the significant and non-significant MPRA effects for several aPCs which reflected the same pattern as Enformer of strongest enrichment in the top percentiles. We also found that these enrichments were strongest for the transcription factor and epigenetics aPCs, while other aPCs like distance from TSS/TES were similarly enriched in both MPRA effects and all other tested variants (Figure 3B inset).

**Figure 3 -.**
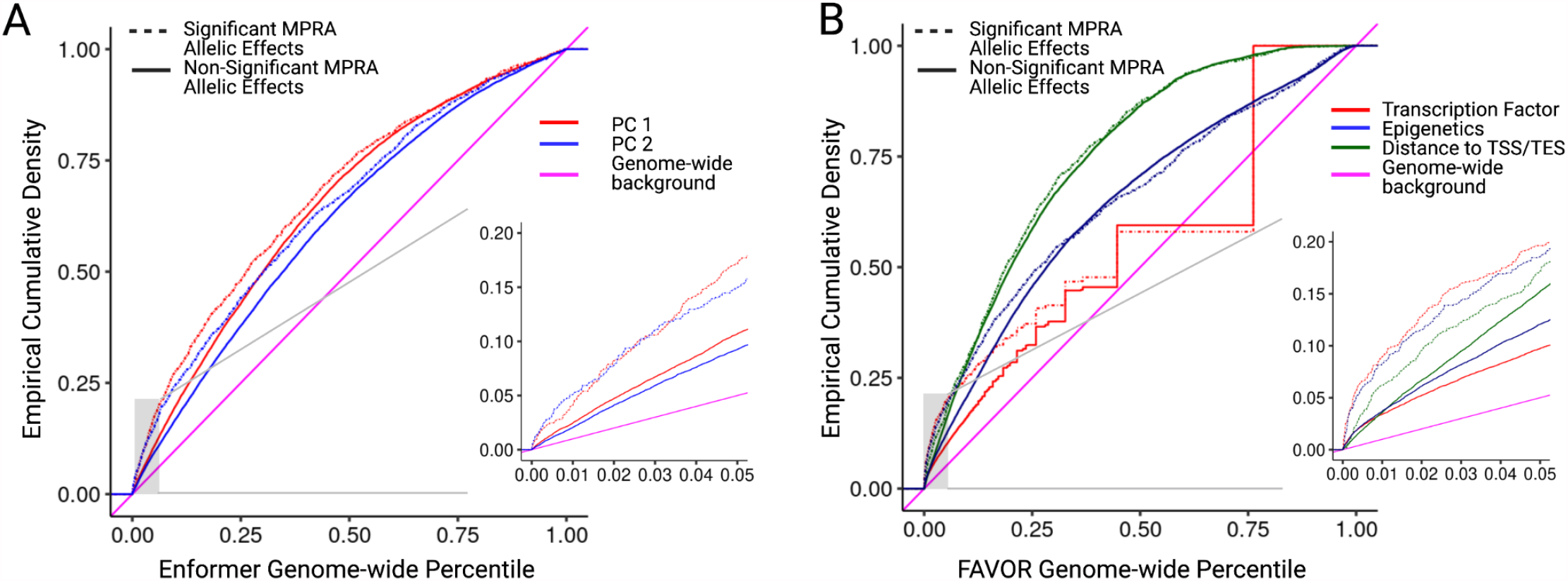
Integrative non-coding variant effect prediction. (A) Empirical cumulative probability distribution of the first and second principal component (PC) scores from Enformer for significant and non-significant MPRA allelic effects; genome-wide percentile computed across all common variants in 1000 Genomes Phase 3. (B) Same as in (A) except showing annotation principle components from FAVOR; genome-wide percentiles computed across all variants in TOPMed Freeze5.

While both predictors could distinguish tested sites in eQTL regions from genomic background, we observed that there was little global performance difference between allelic hits and tested non-hits, except at very high percentiles of Enformer and FAVOR scores (Figure 3A,3B). These results demonstrate that genome-wide non-coding variant effect predictors can easily distinguish regulatory regions near eQTL, but only enrich for causal variation within eQTL at the extreme ends of the genome-wide distribution. This further indicates why functional fine-mapping approaches have not systematically benefitted from non-coding variant effect predictions, while also suggesting that the highest genome-wide percentiles of these integrated scores provide a large source of variants significantly enriched for MPRA allelic effects.

### Multiple causal regulatory variants in high linkage disequilibrium underlie eQTL

In order to fine-map high-confidence causal variants underlying statistical associations, we compared eQTL summary statistics with MPRA allelic effects. Across all loci, 76.7% and 45.6% had at least one such expression effect or allelic effect respectively (Figure 4A). 17.7% of all loci had more than one significant allelic effect, indicating that an appreciable number of individual associations contain multiple regulatory variants (Figure 4B). Even when requiring a very strong expression effect size (absolute log2 effect size > 1.4) in addition to significant expression and allelic adjusted p-values, 6.3% of all eQTL contained multiple active variants.

**Figure 4 -.**
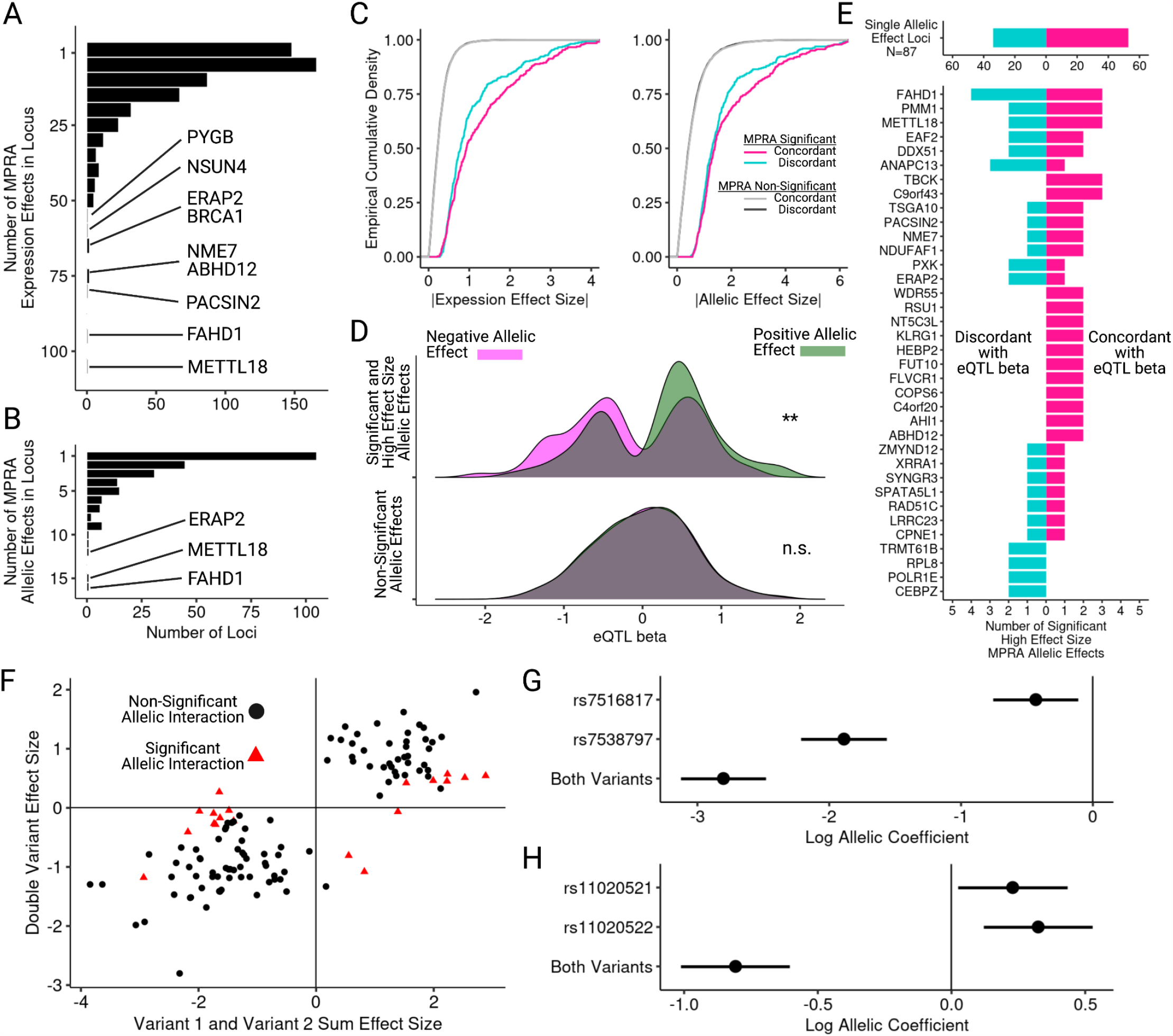
Decomposition of allelic heterogeneity within regulatory loci. (A) Histograms of the number of expression effects per locus with adjusted expression p-value <= 0.05. (B) Same as in (A) except also requiring adjusted allelic p-value <= 0.05. (C) Cumulative distribution of effect size absolute values stratified by concordance; concordance is defined as the sign of the allelic effect size matching the sign of eQTL beta. (D) Distribution of eQTL beta measured in GTEx v8 LCLs for strong MPRA effects (log expression effect size >= 1.5), stratified by MPRA allelic effect direction and significance. (E) Using the same variants as (D), counts of directionally concordant and discordant significant allelic effects across all loci with at least one effect. (F) Comparison of haplotype regression coefficients for variants tested individually or jointly; red points indicate allelic interaction adjusted p-value <= 0.05. (G) Haplotype regression coefficients and standard errors from DESeq2 for two linked variants associated with *FLVCR1* expression and separated by 4 bp. (H) Same as in (G) except for two linked variants associated with *C11orf54* expression and separated by 58 bp. * <0.05, **<0.005, ***<0.0005, ****<5e-5.

The degree to which eQTL are the composite product of multiple causal variants is unknown due to high LD. We assessed whether MPRA allelic effects found within an eQTL were more likely to be concordant with the measured eQTL effect direction than other variants. We found concordant variants had larger expression and allelic effect magnitudes than discordant variants indicating that discordant variants more often have weaker effects (Figure 4C). Across allelic hits with the largest effect sizes (absolute log2 effect size > 1.4), when comparing concordance based on MPRA significance we observed that allelic effects were significantly concordant with eQTL effect direction (Fisher’s exact p-value = 4.75e-3) while effects from non-significant variants were not (p-value = 0.478; Figure 4D). For eGenes regulated by these variants, almost twice as many were completely concordant than discordant when either one or two active variants were present. Taken together, these eGenes provide the strongest examples of regulation by multiple allelic effects in the context of widespread heterogeneity (Figure 4E).

To rule out study-specific effects, we verified that included eQTL effect sizes were highly consistent across multiple studies (Supplementary Figure S4A) (*32, 33*). Indeed, we find the same concordant pattern when using either Geuvadis or GTEx LCL eQTL data (Supplementary Figure S4B). Additionally, to ensure that the observed concordance patterns were not driven by individual eQTL with many concordant variants, we applied a binomial count logistic regression model to test whether concordance proportions were significantly shifted between significant and non-significant MPRA hits while accounting for variation in the total number of tested variants per region. We found that allelic MPRA effects, but not other variants, were significantly more concordant than expected (p-value = 2.85e-3; Figure S4C). Finally, given that concordant variants have broadly larger effect sizes, we evaluated whether the top MPRA variant per eQTL as ranked by allelic effect magnitude was more concordant than other variants. We found that increased concordance persists through the top four ranked variants per eQTL, with the set of third-strongest MPRA hits across all eQTL having a concordance rate of 0.67 (Supplementary Figure S4D). Altogether, these results imply that several eQTL regions contain multiple, concordant, and high effect size allelic MPRA hits and thus may be driven by multiple causal variants.

### Haplotype decomposition identifies allelic regulation that is unlikely to be observed by population sampling

A major advantage of synthetic library design is the ability to separate the effect of extremely proximal variants which are unlikely to be naturally separated by recombination. Our sample set included 2,097 pairs of eVariants that are separated by less than 75 bp. To our knowledge, all previously described allelic interactions of this kind have been reported in *S. cerevisiae* (*19, 20*). For these variants, we extended our initial statistical model to account for all four possible haplotypes at each pair of variants and computed summary statistics for each of the three non-reference haplotypes (Supplementary Figure S5A). We then selected all variants with at least one significant haplotype. Combined, we identified 120 variant pairs (6.15% of all tests) with at least one significant allelic haplotype relative to all-reference sequence (adjusted p-value < 0.05).

Most of the haplotype effects appeared additive (Figure 4F). For example, the haplotype pair rs7516817 and rs7538797 displayed an additive pattern where all three non-reference haplotypes had significant individual effects and the double-variant haplotype displayed a larger effect size than either variant individually in the same direction (Figure 4G). Our linear contrast test also allowed us to identify significant, non-additive interactions between variants (Supplementary Figures S5B-D, Supplementary Table S7). Of the variant pairs with at least one significant haplotype, 19 pairs also had a significant haplotype interaction effect (14.7% of all significant haplotype effects and 0.91% of all tested variant pairs). Significant interactions did not reverse the direction of the individual variant effects completely except in rare cases. For example, the rs11020521 and rs11020522 haplotype pair exhibited an interaction with sign reversal where each variant individually has no effect or is weakly positive, while the joint haplotype is much more significant in the opposite direction (Figure 4H). These results support other synthetic, eQTL and complex trait studies that have identified non-additive regulatory effects (*20, 34–36*) and find that 14.7% of significant haplotype effects (only 0.91% of all tests) have evidence of non-additivity.

### Experimental fine-mapping of complex trait associations

In order to identify loci with shared genetic architecture between our eQTL and human traits, we retrieved all genes that were tested in our dataset and had both a significant MPRA hit and one or more significant potential colocalizations between LCL eQTL and a curated GWAS set (*37, 38*). Out of 744 eGenes, 5.51% both colocalized with at least one trait and contained at least one significant allelic effect. Notably, the majority of these colocalizations contained more than one such effect (71.9% of colocalizations and 82.9% of eGenes), with some loci containing as many as 13 active variants (Figure 5A). This suggests that the default assumption of one causal variant, often used in approaches for fine-mapping or colocalization of GWAS loci, does not reflect causal variant biology at many regulatory regions. Traits with high-confidence co-localization were diverse, including blood-cell traits like *ZC2HC1A*/Lymphocyte Count or *PACSIN2*/Platelet Count, as well as significantly more complex traits like *GNA12*/Height (Supplementary Figure S6).

**Figure 5 -.**
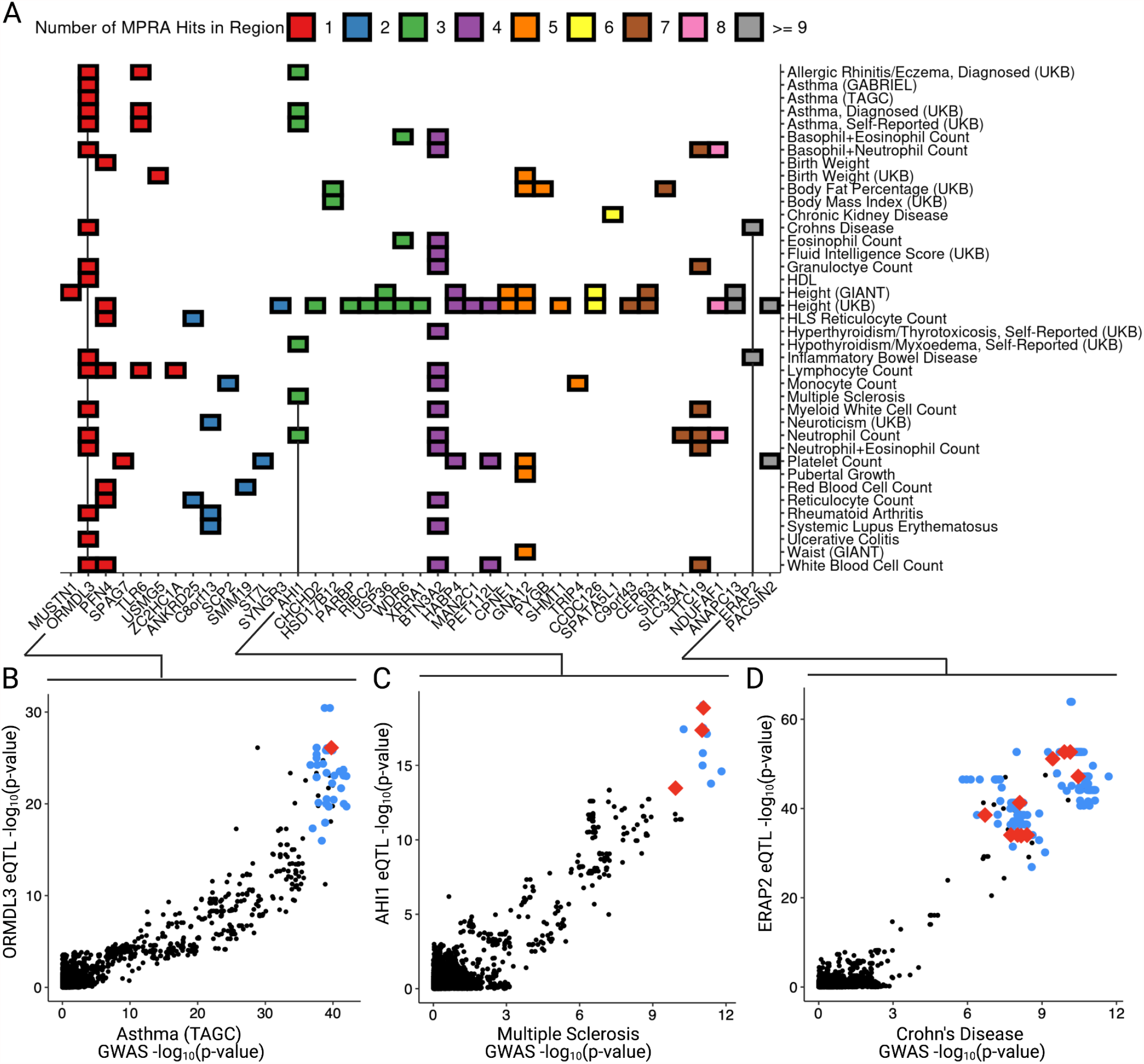
Resolving complex trait associations with multiple causal variants. (A) Heatmap of significant colocalizations between eQTL loci and selected GWAS; color indicates the number of significant MPRA allelic effects within the colocalized region including both single and double variant effects. (B) Comparison of variant p-values for association with Asthma and *ORMDL3* expression in GTEx v8 LCLs; red and blue points indicate significant and non-significant MPRA allelic effects respectively, black points indicate untested variants. (C) Same as in (B) for associations with Multiple Sclerosis and *AHI1* expression. (D) Same as in (B) and (C) for associations with Crohn’s Disease and *ERAP2* expression which was also colocalized with Inflammatory Bowel Disease (Supplemental Figure 5A). All GWAS and eQTL colocalizations are retrieved from (*38*), and lead variants were always required to be genome-wide significant even if colocalization probability was high.

One colocalization that was consistent with the single causal variant assumption was *ORMDL3*, which has been extensively studied as an asthma susceptibility gene. Our GWAS set included two UK Biobank datasets (doctor diagnosed and self-reported) and two independent asthma studies. We observed colocalization across all of them with *ORMDL3* expression and identified a single significant allelic hit, rs12950743, which was highly significant but not the most significant in all incorporated association studies (Figure 5B). This variant has been previously associated with CpG methylation and chromatin accessibility in several studies, but linkage precluded single-variant analysis (*39, 40*). Interestingly, in a different study this variant was associated with asthma and specifically identified as “mechanistically likely” although *ORMDL3* was not identified as its target gene (*41*). Taken together, these results provide new empirical evidence supporting rs12950743 as a causal variant driving both *ORMDL3* expression and asthma susceptibility.

In contrast, a different colocalization that included three active variants was *AHI1*, which is a well-characterized gene containing variants strongly associated with Multiple Sclerosis (MS) (Figure 5C) (*42–44*). This region has been the subject of multiple functional studies and contains a strong eQTL and colocalization signal in LCLs and many other blood cell types; however, its causal variant(s) has not yet been resolved. We identified rs6908428, rs9399148, and rs761357 as displaying significant allelic effects. The first two of these variants have been highlighted by annotation overlap in variant sets from prior associative studies that performed extensive characterization of the role of *AHI1* regulation in MS pathology and CD4+ T-cell pathology, but which were severely limited by linkage across the risk haplotype (*43*). Our results suggest that both of these variants, along with rs761357, may regulate *AHI1* expression and thus contribute to MS risk.

One of the most complex multi-variant colocalizations was identified at the *ERAP2* locus, which is an aminopeptidase that has been functionally implicated in both Inflammatory Bowel Disease and Crohn’s Disease and for which we observe very strong eQTL-GWAS colocalization (*45–47*). We detected thirteen active variants which span a strongly linked haplotype. While both eQTL and GWAS data suggest a single top SNP, that top SNP differs between eQTL and GWAS and neither demonstrated any significant MPRA effect (Figure 5D, Supplementary Figure 6A). Prior work has shown that a common splice variant in *ERAP2* results in nonsense-mediated decay (NMD) and allele-specific expression, which can cause an eQTL signal (*48*).

However, the haplotype with this variant contains hundreds of other linked variants with significant population variation and harbors a second conditional *ERAP2* eQTL in GTEx v8 LCLs. Moreover, several prior studies have demonstrated associations between *ERAP2* expression and cis-variation independently of NMD, although linkage limited the resolution of functional studies (*33, 45, 49, 50*). Taken together, these results suggest that *ERAP2* is regulated by a complex multi-allelic structure which operates via both gene expression and splicing.

## DISCUSSION

Linkage disequilibrium is a major barrier to identifying causal variants in genetic association studies, particularly for the vast majority of significant associations which are non-coding or for molecular phenotypes like gene expression with limited sample sizes. Furthermore, functional annotations derived without respect to genetic variation are useful to prioritize variant subsets but are often inconclusive, unattainable, or unknown (*2*). In this study, we demonstrate that MPRAs provide a scalable platform to separate and map the regulatory activities of complex trait-associated natural genetic variants, and highlight the limitations of existing approaches to variant interpretation and computational fine-mapping. Across positional annotations and variant scores, we frequently observed that both our significant and non-significant allelic MPRA variants were highly shifted relative to the corresponding genome-wide distributions. While unsurprising since variants were predominantly selected from known eQTL, this effect demonstrates how functional predictions may be capable of readily distinguishing eQTL regions from genomic background while struggling to discriminate regulatory activity between highly linked and jointly significant variants within the same region.

As many statistical fine-mapping approaches rely on functional genomic annotations, our study allowed us to investigate the proportion of predicted binding events that exhibit MPRA transcriptional effects. We found that while our significant allelic MPRA effects were both enriched in and much more concordant with properties like transcription factor binding and chromatin accessibility than non-significant effects, most variants with significant binding scores did not show either an expression or allelic effect in our assay. This may be due to the use of an episomal assay (*51*), technical noise, the absence of cell-type and chromatin context dependencies, or may reflect current assumptions that an appreciable proportion of variants that display allelic imbalance in binding or accessibility have little downstream effect on transcription (*52, 53*). Among others, members of the ETS-domain family including ELK1, ELF1, and ETV1 showed consistent enrichment in MPRA hits in both positional overlap and direction of variant effect.

Finally, we found that multiple causal variants are often found under many eQTL and GWAS loci. We identified that at least 17.7% of eQTL and 71.9% of their colocalizations had more than one allelic hit. We further observed that most combinations of variants exhibited additive effects with 14.7% of active haplotype pairs (0.91% of all pairs) exhibiting non-additivity. Using these combined data, we demonstrate the power of MPRA-based experimental fine-mapping and report likely causal variants underlying hundreds of molecular and complex trait phenotypes, including a single variant underlying *ORMDL3*/Asthma, three variants underlying *AHI1*/Multiple Sclerosis, and up to thirteen variants underlying *ERAP2*/Crohn’s Disease/Inflammatory Bowel Disease.

## METHODS

### cis-eQTL identification in 1000 Genomes population groups

Whole genome sequences and gene expression data were obtained from unrelated individuals from the following populations: Utah residents with Northern and Western European ancestry (CEU); Han Chinese from Beijing, China (CHB); Gujarati Indians in Houston, Texas, USA (GIH); Japanese in Tokyo, Japan (JPT); Luhya in Webuye, Kenya (LWK); Mexican ancestry in Los Angeles, California, USA (MEX); and Yoruba in Ibadan, Nigeria (YRI). Whole genome sequences were from the 1000 Genomes Project Phase 3 (v5).

For analysis of *cis*-eQTL in individuals sequenced in the 1000 Genomes Project Phase 3, 69 individuals from each of CEU, CHB, GIH, JPT, LWK and YRI were selected where each individual had gene expression data previously available (*54*). Variant sites were filtered for a minor allele frequency of at least 1% across all six populations, and at least three non-reference alleles in the population were required to be tested for association to gene expression. Variant positions were coded as 0, 1 or 2 non-reference or alternate alleles, and gene expression was quantified and normalized as previously described (*54*). We used linear regression to test variants within 1 Mb of the transcription start site within each population. We further meta-analyzed *cis-*eQTL between populations using Fisher’s combined *p*-value. Across eQTL, all variants with the most significant p-value in the single population CEU analysis or meta-analysis were considered as lead variants for subsequent MPRA design. We identified 744 such eGenes and lead variants (Supplementary Table S1).

### Lead variant selection and LD expansion

After lead variants were identified across all eQTL, we computed linkage disequilibrium estimates within our dataset between each lead variant and all other significant tested variants in the same *cis*-window. All variants with an r^2^ >= 0.85 were selected for inclusion, resulting in a set of 30,129 variants. For all variants, if another variant occurred within 75bp we included all possible haplotypes centered at both variants, expanding our test set to 41,964 allelic pairs. Additionally, we constructed two smaller variant sets. First, we identified 2,363 variants without any significant eQTL associations in our dataset to any gene (all nominal p-values > 0.25) but were derived from the same *cis*-windows. Second, as a control set, we extracted the top 2,373 variants from (*14*) ranked by combined allelic FDR. This set included the 842 variants identified as emVars in (*14*), as well as 1,531 additional variants (Supplementary Table S2).

### Oligonucleotide library sequence design and synthesis

To construct synthesis-ready sequences, a 150bp window was recovered from the human genome centered at each variant. If multiple variants were present in the same window, we designed all possible haplotypes centered at each variant. Additional sequences were added to the 5’ (ACTGGCCGCTTGACG-) and 3’ (-CGCAGGAGCCGCAGTG) ends of each fragment and synthesized in a 100K pool by Agilent Technologies (Supplementary Table S2). Upon receipt, oligos were resuspended at 2 ng/ul in TE buffer, aliquoted and stored at −20C.

### Reporter plasmid library construction

To accommodate a large number of variants, we implemented the random barcoding procedure described in (*14*) and attached random 20bp barcodes to each oligo fragment with emulsion PCR in 96 parallel reactions using MPRA_Forward and MPRA_20N_Reverse primers (Supplementary Table S9). Following cloning into pMPRA1 (Addgene #49349) (*8*), the library was electroporated into 10-beta electrocompetent *E. coli* in six parallel transformations using a BioRad Gene Pulser. Cultures were independently recovered in 1mL of SOC media, followed by independent outgrowth in 20 mL each of LB media with 100ug/mL carbenicillin for 14 hours. After growth, cultures were pooled and purified by midiprep according to standard protocols. Upon sequencing, we detected approximately 12.2 million barcodes with >98% of designed oligos represented at a median of 121 sequences per oligo in the pool.

To prepare our final library, a minimal promoter attached to GFP (minP-GFP) amplicon was ordered as a gBlock from Integrated DNA Technologies (IDT), amplified, and inserted into the plasmid library by Gibson assembly as in (*14*). The final plasmid pool was electroporated as before except in nine parallel transformations followed by independent recovery in 250 mL LB with carbenicillin. After outgrowth all cultures were pooled, purified by Qiagen gigaprep, resuspended in water and quantified by NanoDrop.

### Mammalian cell culture and transfection

GM12878 cells were cultured in RPMI supplemented with 15% FBS and 1% penicillin/streptomycin maintaining a density of 0.2-1.0×10^6 cells/mL at 37C and 5% CO2. For each replicate, 5×10^8 cells were collected by centrifugation, washed with PBS, and resuspended in 5 mL RPMI containing 500 ug of plasmid library. Each replicate was transfected by electroporation with the Neon system using the following program: [1200V, 20ms, 3 pulses]. Electroporations were performed in 100 ul volumes, and each tip was used five times while each electroporation buffer tube was used ten times. After electroporation, cells were immediately recovered in 500 mL of RPMI with 15% FBS and no antibiotics for 24 hours at 37C.

### Reporter mRNA isolation and normalization

After recovery, total RNA was isolated using 4 mL Trizol per replicate following the standard protocol. All RNA was used as input for Turbo DNase digestion according to the manufacturer’s protocol except scaled to 500 uL per reaction. Following digestion, GFP mRNA was recovered using biotinylated antisense oligonucleotides targeting the GFP mRNA using probes GFP_Biotin_[1-3] obtained from IDT exactly as described in (*14*). Following extraction, recovered mRNA was DNase I digested again except in a 100 µl reaction. cDNA was generated by reverse transcription with SuperScript III according to the manufacturer’s protocol with a gene specific primer (MPRA_Amp2Sc_R, Supplementary Table S9) and purified by SPRI into 30 µl of water.

### Oligo-barcode mapping and barcode sequencing

For sequencing of barcodes from plasmid or cDNA, the following PCR reaction was performed with each sample: [25 µl Q5 NEBNext Master Mix, 2.5 µl MPRA_Illumina_GFP_Forward, 2.5 µl Illumina_Universal_Adapter 10 µl water, 10 µl sample] and cycled for [95C 2 minutes, [95C 20 seconds, 65C 15 seconds, 72C 30 seconds] x 12, 72C 2 minutes]. Before amplification, all samples were first normalized by qPCR as described in (*14*), using the same reaction except in 10 µl and with the addition of Sybr Green I (Supplementary Table S9). All samples were 1.8X SPRI purified into 30 µl water for indexing.

For sequencing of oligo-barcode pairs, four parallel PCR reactions were performed with the following composition: [100 µl Q5 NEBNext MasterMix, 10 µl MPRA_Amp2Sa_Illumina_Forward, 10 µl Illumina_Universal_Adapter, 78 µl water, 2 µl plasmid library (100 ng/µl)] and cycled for [95C 2 minutes, [95C 20 seconds, 62C 15 seconds, 72C 30 seconds] x 6, 72C 2 minutes]. Reactions were pooled and 1.8X SPRI purified into 30 µl of water for indexing (Supplementary Table S9).

For indexing and sequencing of all libraries, multiplex adapters were added using the following PCR reaction: [50 µl Q5 NEBNext Master Mix, 5 µl Illumina_Multiplex_[1-10], 5 µl Illumina_Universal_Adapter, 10 µl water, 30 µl sample] and cycled for [95C 2 minutes, [95C 20 seconds, 65C 30 seconds, 72C 30 seconds] x 6, 72C 2 minutes]. After amplification, samples were 1.5X SPRI purified, checked by Agilent Bioanalyzer, and pooled for sequencing on an Illumina NovaSeq 6000.

All primer sequences are included in Supplementary Table S9, and in all described reactions primer concentrations were 10 µM for a final concentration of 0.5 µM in reaction. Oligo-barcode pairs were sequenced on NextSeq 500 and NovaSeq 6000 instruments, and barcodes were sequenced using the NovaSeq 6000.

### Sequencing data analysis

To map barcodes to their parent oligo sequences, we used FLASH to merge paired-end reads derived from the merged NextSeq and NovaSeq data (*55*). Random barcodes were extracted from each merged fragment and appended to the read name. We used STAR v2.7.1a to align merged reads against a reference index created from the designed library sequences (*56*). After filtering reads that did not map uniquely to a designed sequence or which had low quality alignment scores (< 100), we extracted the resulting barcode-oligo pairs and removed any sequences detected on multiple oligos.

To quantify oligo-level counts from barcodes, we used Bartender v1.1 on each sample individually to obtain barcode clusters to correct for sequencing errors relative to our known barcode list (*57*). After clustering we computed oligo counts by mapping each barcode to its parent oligo requiring an exact match, and summed all barcode counts within each oligo. The final oligo count matrix contained measurements for 46,699 allelic pairs (Supplementary Table S3).

### Statistical modeling and inference of MPRA regulatory effects

To account for variation due both sequencing depth and allelic ratios, we applied a nested fixed model using DESeq2 described for high-depth allele-specific expression analysis that accounts for the intrinsically paired allelic design (*22, 58*). We required a raw count mean >= 150 across all samples to remove very low count oligos, and then used the following nested model:

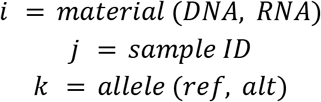

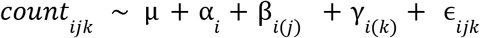

Here, α describes the aggregate allele-independent effect observed in RNA compared to DNA, while *β* and γ describe sample-specific and allele-specific effects respectively, nested within either RNA or DNA as denoted by the subscript parentheses. Using a linear contrast, we extracted the global RNA vs DNA average effect as “expression” effects, and significantly non-zero nested allelic terms indicate “allelic” effects. In this way, we obtained summary statistics that correspond well to the fold change and skew values reported in (*14*), respectively, but do so with an integrated model that does not require separate inference of allelic effects after expression effects (Supplementary Figure S2).

After fitting, we tested for expression effects using a Wald test on α and tested for allele effects using a linear contrast between DNA and RNA levels of γ to test the null hypothesis that the nested allelic coefficients were equal. For both sets of summary statistics, p-values were adjusted for multiple testing using the Benjamini-Hochberg procedure. Our final dataset included 30,523 tests for both expression and allelic effects for single variants (Supplementary Table S4) and 2,097 tests for haplotype effects (with 6,291 summary statistics as there are three non-reference haplotypes per pair) between two variants (Supplementary Table S7).

### Functional genomic data and annotation

To annotate tested variants with uniform chromatin immunoprecipitation (ChIP-seq) data, we downloaded all tracks from the ReMAP 2020 database derived from GM12878 cells which resulted in usable data for 160 transcription factors (*23, 24*). As this set does not include human histone modifications, we separately downloaded all ENCODE histone tracks derived from GM12878 cells which resulted in 15 datasets of which 11 had sufficient overlap for testing, spanning 8 marks (Supplementary Table S6). To obtain RNA-seq measurements of transcription factors themselves, we downloaded GM12878 RNA-seq data quantified in transcripts per million (TPM) from ENCODE. All odds ratios were computed as Fishers’ exact tests (Supplementary Table S5).

We analyzed SNP-SELEX data by downloading annotations provided by for significant scores across all common variants in 1000 Genome Phase 3 (*26, 59*). Variants were matched based on chromosome, position, and alleles but not by strand. Concordance was assessed using the sign of the ‘deltaSVM_score’ field from SNP-SELEX annotations and the sign of the MPRA allelic effect size.

For positional chromatin accessibility enrichments, we downloaded cell-type specificity and footprint identity annotations from (*25*) and overlapped with tested MPRA variant coordinates using bedtools. We imposed no additional filtering or thresholding on the interval sets provided. Separately, for caQTL and allelic imbalance analyses, we obtained data from (*28*) and from (*29*) which both characterize the effects of genetic variation on chromatin accessibility. We required all variants to have at least a nominal p-value of < 0.50 for allelic imbalance. We then assessed concordance by subtracting 0.5 from the allelic proportions and comparing the sign with the MPRA allelic effect direction. For caQTLs from (*29*), we evaluated overlap based on chromosome, position, alleles, and the number of populations demonstrating the caQTL effect and required concordance to consider an overlap valid.

### Integrated variant effect predictors

Enformer variant effect score principal components were obtained from (*30*) and matched to tested variants based on chromosome, position, and alleles. To compute the genome-wide background, we downloaded all provided PC scores across common variants on chromosome 22 in 1000 Genomes and computed reference quantiles. PC scores on tested variants were then scaled to this distribution and used for comparisons based on MPRA effect significance.

FAVOR scores for annotation principal components (aPCs) are provided raw and Phred scaled, so we downloaded aPC scores for only tested variants and recovered their genome-wide percentile using the inverse Phred transformation (*31*). Unlike for Enformer scores, we did not directly compute a genome-wide distribution but rely on the rank-scaling of reported FAVOR Phred scores across all variants in TOPMed Freeze5.

### eQTL integration and concordance analysis

We designed our library based on significant eQTL variants from 1000 Genomes populations. To ensure the extensibility of our data, we downloaded eQTL data for LCLs from GTEx v8 and compared estimated effect sizes between all matching tests (same variant and eGene) (*33*). When considering allelic MPRA effects, we required that effect size direction be consistent and nominally significant (unadjusted p-value < 0.05) in both eQTL studies. For all concordance analysis, the sign of a given eQTL effect size was compared with the sign of the corresponding allelic MPRA effect. To identify eGenes with the strongest evidence of multi-variant regulation, we required an absolute expression log2 fold change >= 1.4 as well as adjusted expression and allelic p-values <= 0.05.

### Haplotype interaction modeling

By design, our MPRA library includes oligo sequences with all possible variant combinations when multiple variants occur within an oligo-length window. To effectively utilize all measurements relevant to each site, we performed a separate analysis of all variant pairs which were located within 75bp of each other and for which we had oligo counts corresponding to all four haplotypes (Supplementary Figure S5A). We implemented this model in DESeq2 as for the general single-variant analysis, except requiring a raw count mean of 400 across all samples and adding an additional component to the nesting structure indicating the oligo position:

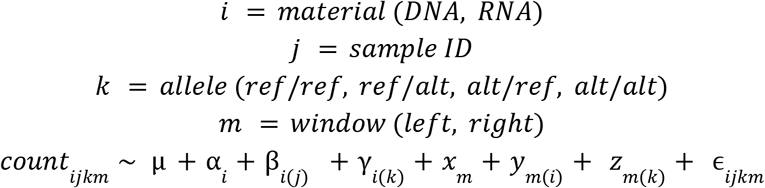

There are three additional terms *x,y*, and *z* that correspond to the overall effect of the window and nested effects within each window for material and allele respectively. We used the same approach as in the single variant analysis, since we aimed to identify variants without significant positional or window-dependent effects, except with three allelic comparisons (each non-reference haplotype to all-reference) instead of one. We applied a linear contrast to obtain global RNA vs DNA effects as well as tested individual terms for significant allelic effects. We had sufficient coverage to test 2097 variant pairs, corresponding to 16776 oligo sequences (4 haplotypes in each of 2 windows). We also applied a linear contrast to test for evidence of non-additive interactions between variants, meaning that the estimated effect of the joint haplotype was significantly different than the sum of the effects of the component variants individually (Supplementary Table S7).

### GWAS co-localization and fine-mapping

Since most association signals were consistent between our initial eQTL data and GTEx, we used data presented in (*38*) within GTEx v8 to identify significant colocalizations between LCL eQTL and 114 curated GWAS covering a range of traits and complex diseases, including 45 GWAS derived from the UK Biobank. For all GWAS, we downloaded transcriptome-wide colocalization results with GTEx v8 LCLs computed using coloc with enloc priors (*37, 38, 60*). We considered colocalizations that overlapped at least one significant allelic effect and for which PP3 + PP4 >= 0.5 and PP4 >= 0.1 where PP indicates a posterior probability within the coloc framework, in order to ensure substantial signal overlap while allowing for differences in lead variants (Supplementary Table S8).

## Code and data availability

All summary statistics and figure generation analyses are available as Jupyter notebooks at https://github.com/nsabell/mpra-v2. Sequencing data are available through the Gene Expression Omnibus under accession GSE174534. Processed oligo counts are at both locations.

## Declaration of Interests

SBM is on the SAB of MyOme Inc. All other authors report no competing interests.

## Acknowledgements

We thank members of the Montgomery lab for general guidance and feedback on this work and members of the Bassik lab for experimental advice. We also thank Ryan Tewhey and Michael Love for experimental and statistical modeling advice, respectively. NSA is supported by the Stanford Department of Genetics T32 training grant and the Joint Institute for Metrology in Biology (JIMB) training program. SBM is supported by National Institutes of Health grants R01AG066490, R01MH125244, U01HG009431 (ENCODE), R01HL142015 (TOPMed) and R01HG008150 (NoVa). This work in-part used supercomputing resources provided by the Stanford Genetics Bioinformatics Service Center, supported by National Institutes of Health S10 Instrumentation Grant S10OD023452.

## SUPPLEMENTARY FIGURES

**Supplementary Figure 1 -.**
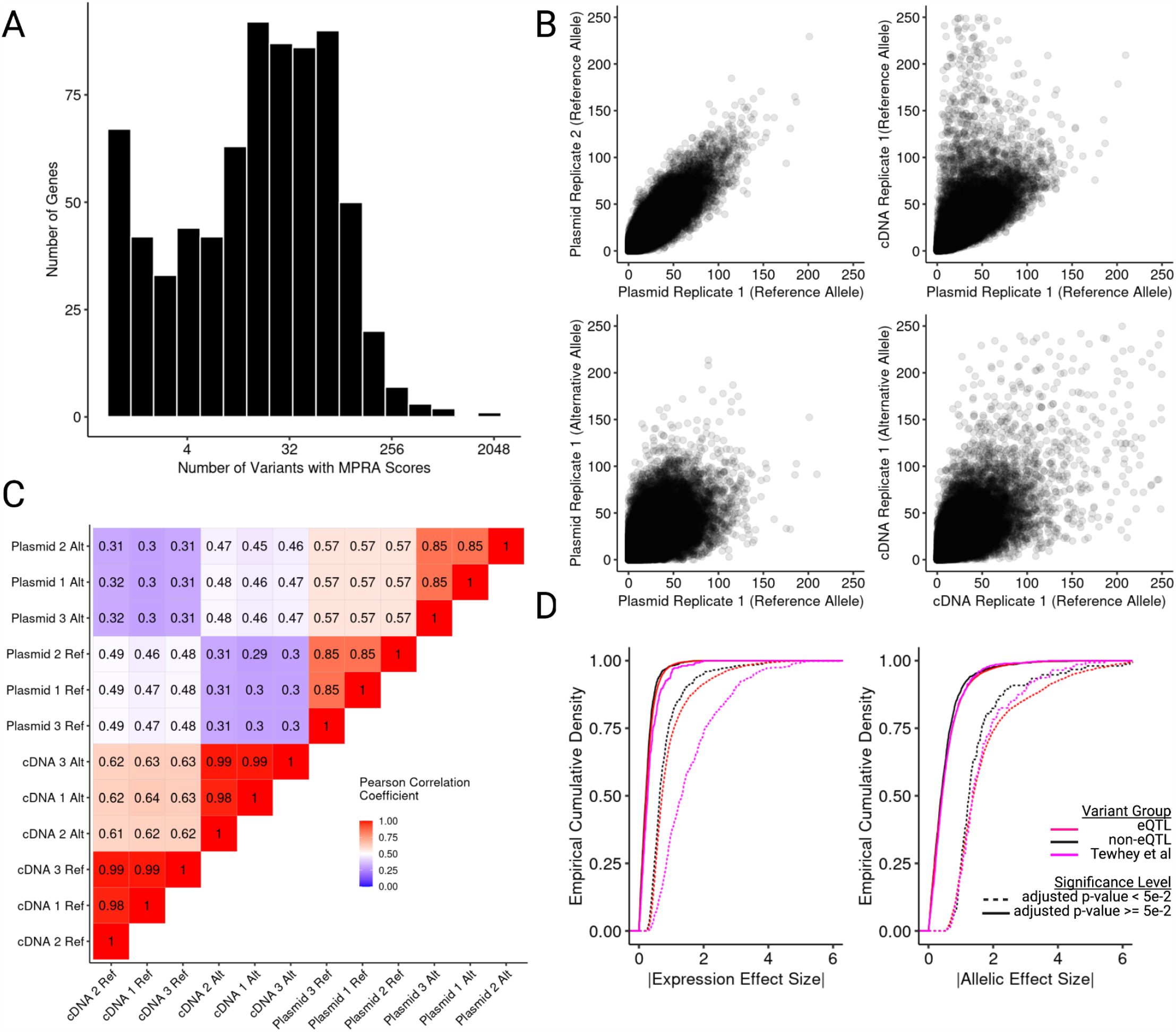
MPRA library design and descriptive statistics. (A) Distribution of the number of variants per eQTL region for which expression and allelic effects were computed and tested. (B) Example pairwise comparisons between plasmid replicates (upper left), one plasmid and one cDNA replicate (upper right), reference and alternative allele counts from the same plasmid replicate (lower left), and reference and alternative allele counts from the same cDNA replicate (lower right). (C) Clustered Pearson correlation heatmap of oligo counts separated by reference and alternative allele for each replicate. (D) Empirical cumulative densities of absolute effect sizes stratified by significance and variant set for expression and allelic effects.

**Supplementary Figure 2 -.**
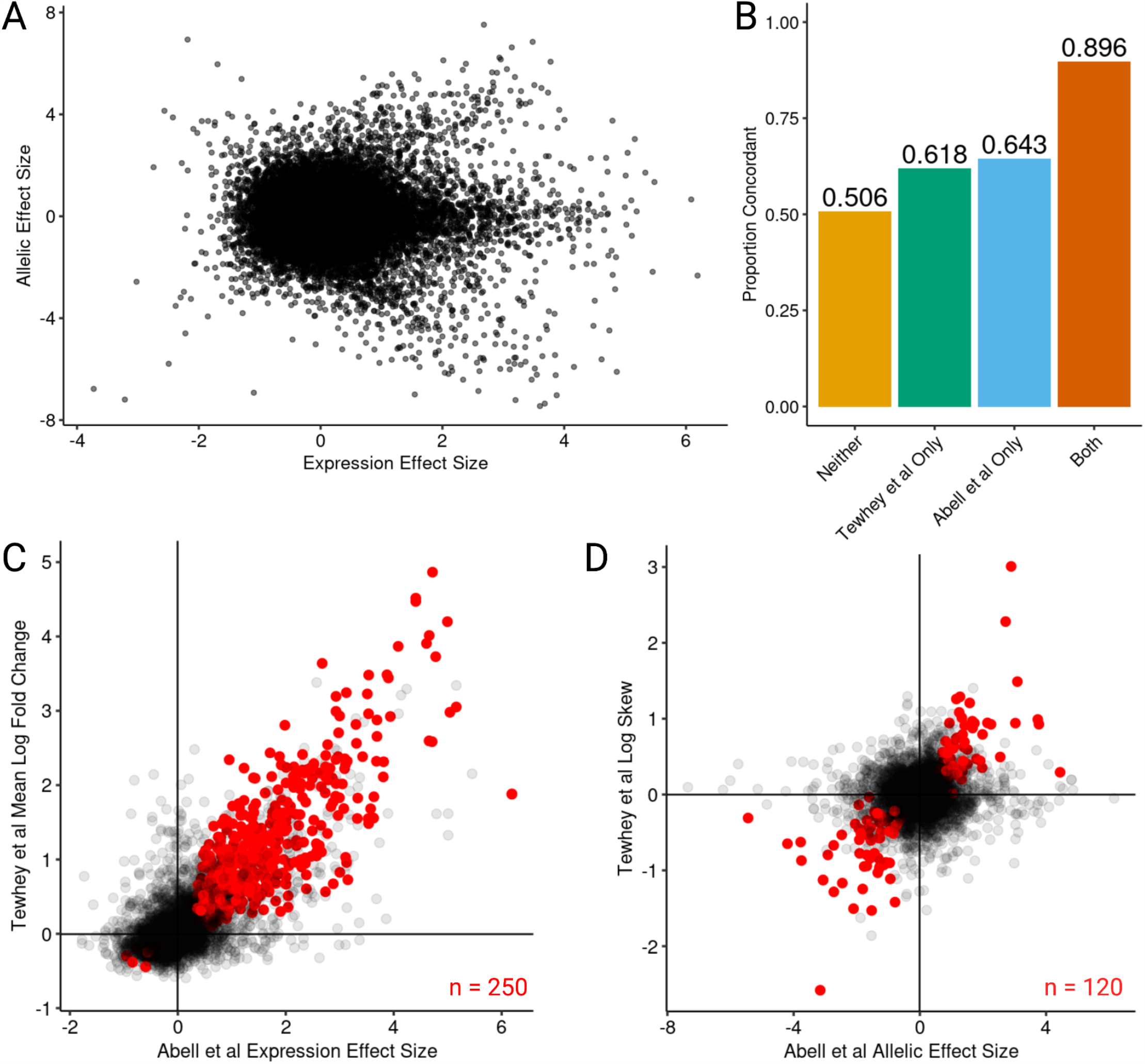
Replication of Tewhey *et al* expression-modulating variants. (A) Expression and allelic effect sizes across all tested variants. (B) Proportion of variants where the sign of the effect size is the same in (*14*) and this study stratified by whether a variant was significant in one, both, or neither dataset. (C) Comparison of expression effect size in this study and the mean log fold change reported in (*14*) for all shared variants (black) and the subset of variants identified as highly reproducible (red). (D) Same as in (C) except showing the relationship between allelic effect sizes from this study and the log skew reported in (*14*).

**Supplementary Figure 3 -.**
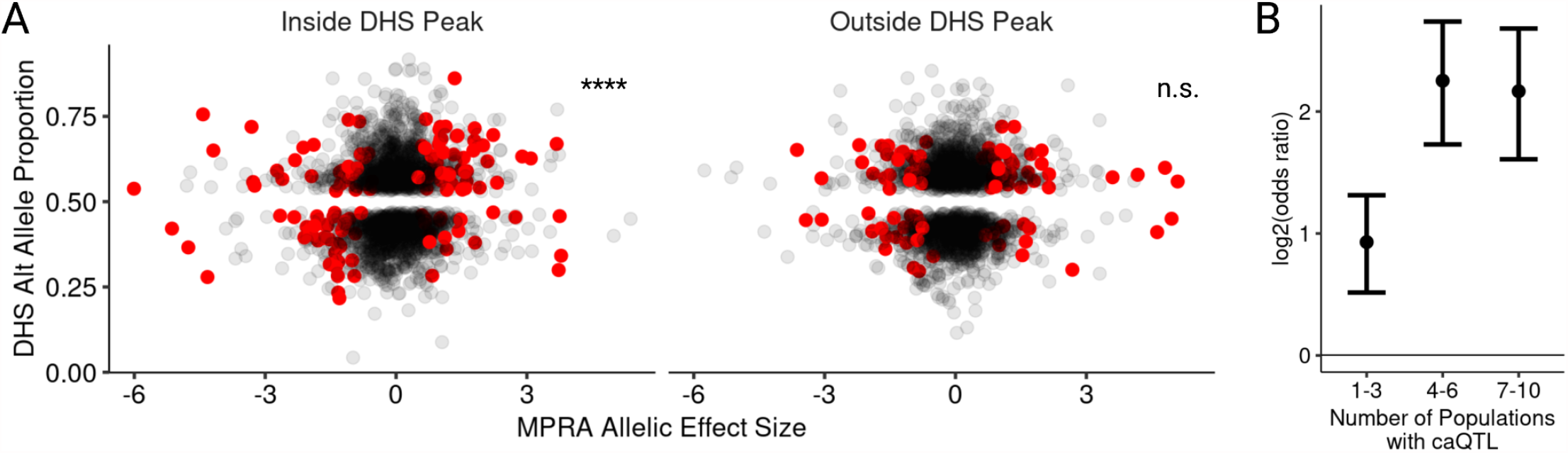
Comparing chromatin accessibility QTLs with allelic MPRA effects. (A) Comparison of MPRA allelic effect size and DHS alternate allele proportion; red points are significant MPRA and DHS-skewed variants, black points are all other variants; only variants with nominal allelic imbalance p-value <= 0.25 were included. (B) Odds ratios and 95% confidence intervals for enrichment of significant caQTLs from (*29*) within significant MPRA allelic effects and varying levels of population sharing. * <0.05, **<0.005, ***<0.0005, ****<5e-5.

**Supplementary Figure 4 -.**
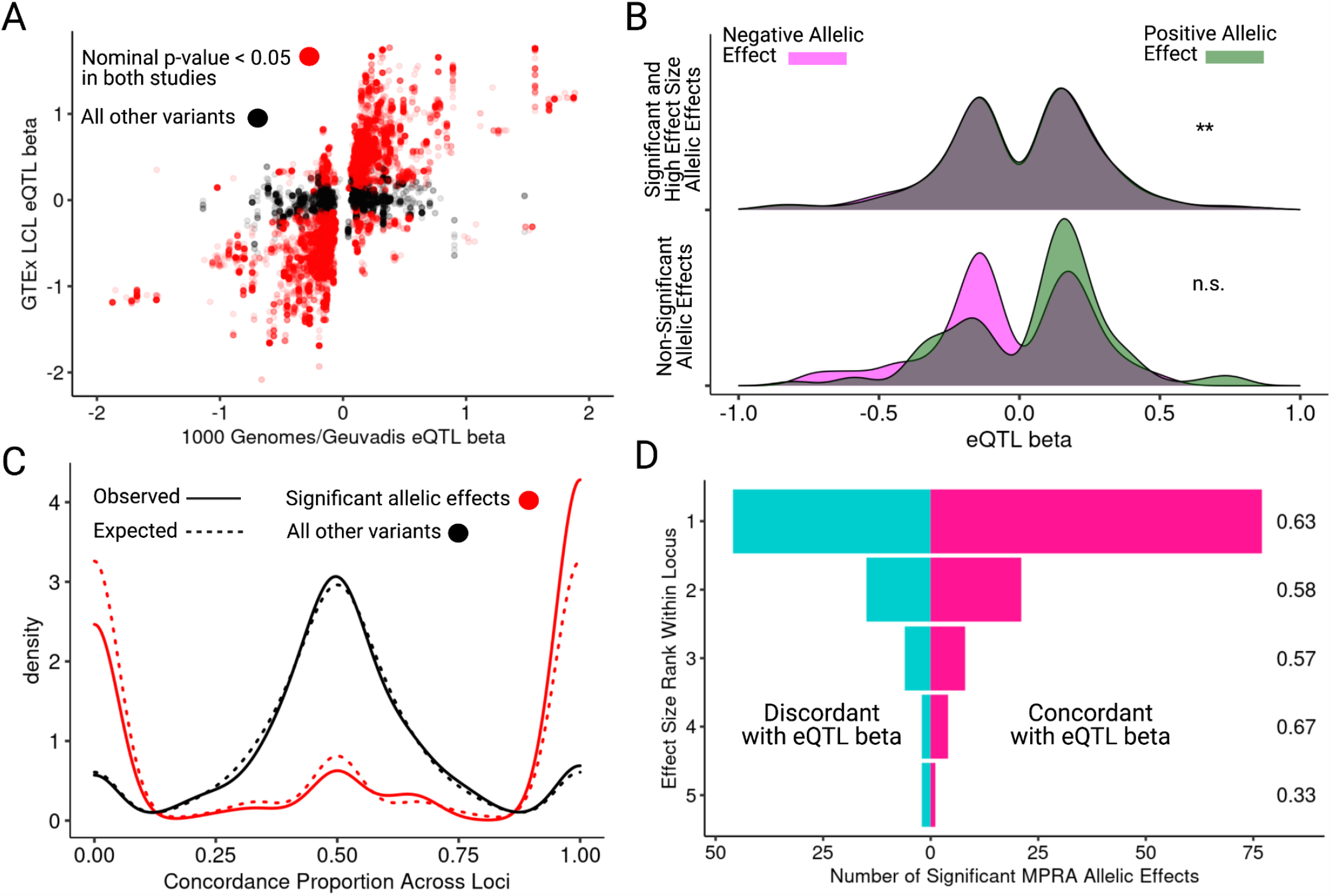
eQTL replication and allelic heterogeneity. (A) Comparison of eQTL effect sizes computed in either Geuvadis (x-axis) or GTEx LCLs (y-axis); red denotes shared nominal significance in both datasets. (B) Distribution of eQTL beta in 1000 Genomes/Geuvadis LCL eQTL stratified by MPRA allelic effect direction and significance. (C) Distribution of observed and expected concordance distributions across all loci for significant and non-significant allelic effects, taking into account differences in the number of tested variants per locus; significance of shift in observed probabilities is tested by binomial regression (p-value = 3.21e-3). (D) Number of significant concordant and discordant variants stratified by allelic effect size rank within locus for variants in Figure 4D and 4E; values to the right indicate concordance proportions. * <0.05, **<0.005, ***<0.0005, ****<5e-5.

**Supplementary Figure 5 -.**
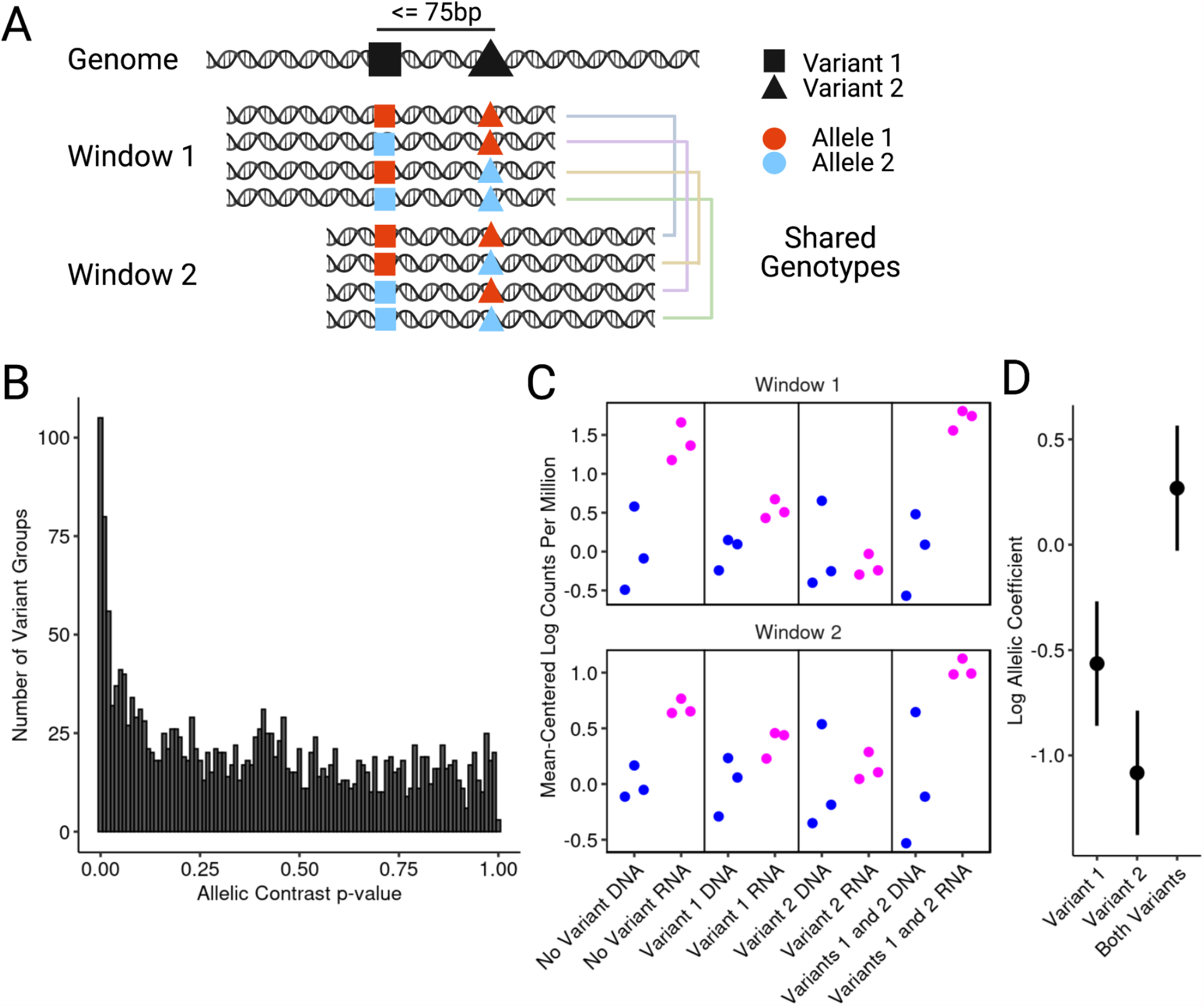
Haplotype decomposition of extremely proximal variation. (A) Motivation for a separate haplotype regression model to take into account repeated inclusion of identical genotypes with shifted windows for very proximal variant pairs. (B) Unadjusted p-value distribution of interaction tests across 2049 haplotype groups; after Benjamini-Hochberg adjustment 37 variant groups had a significant interaction adjusted p-value <= 0.05. (C) Example locus showing log2 counts per million for each measurement included in a haplotype regression; for each pane, the DNA mean (blue points) was subtracted from all points. (D) Haplotype regression coefficients and standard errors from DESeq2 for the haplotype group shown in (C).

**Supplementary Figure 6 -.**
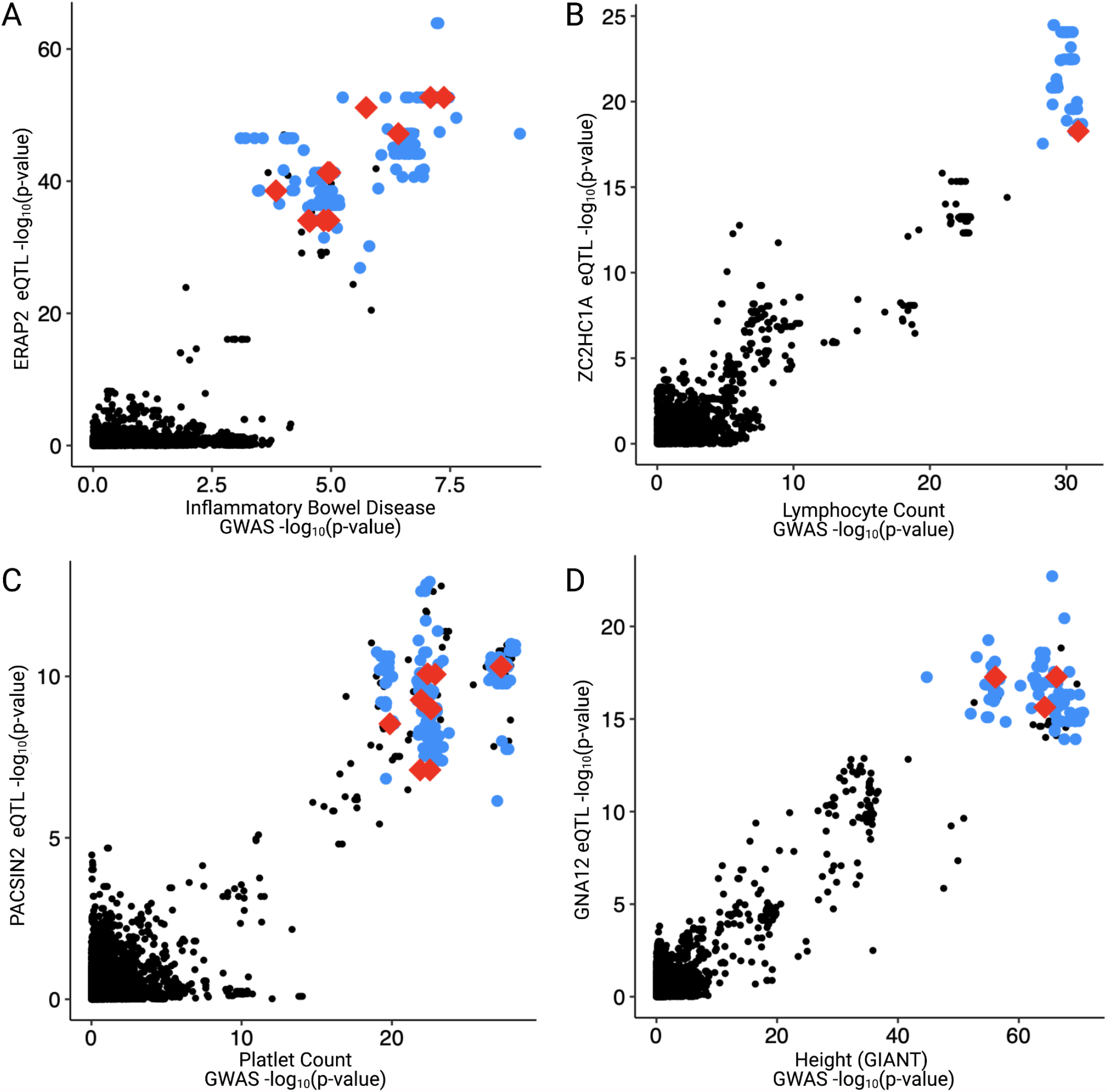
Additional selected GWAS-eQTL colocalizations with varying degrees of allelic complexity. Additional GWAS-eQTL colocalizations containing various numbers of putative causal variants. For all comparisons, both eQTL and GWAS peaks contain a genome-wide significant signal. (A) *ERAP2-*Inflammatory Bowel Disease (nine active variants). (B) *ZC2HC1A*-Lymphocyte Count (one active variant). (C) *PACSIN2*-Platelet Count (ten active variants). (D) *GNA12*-Height (three active variants).

## SUPPLEMENTARY TABLES

Table S1 - Lead variant eQTL summary statistics

Table S2 - Library design sequences and IDs

Table S3 - Oligo count matrix

Table S4 - DESeq2 summary statistics

Table S5 - Odds ratios/intervals for all enrichment tests

Table S6 - Table with descriptions and URLs of all external datasets

Table S7 - DESeq2 summary statistics, haplotype analysis

Table S8 - Colocalization posterior probabilities

Table S9 - Primer and cloning sequences

